# A Dystroglycan–Laminin–Integrin axis controls cell basal geometry remodeling in the developing *Drosophila* retina

**DOI:** 10.1101/2022.11.28.518187

**Authors:** Rhian F. Walther, Courtney Lancaster, Jemima J. Burden, Franck Pichaud

## Abstract

Cell shape remodeling is a principal driver of epithelial tissue morphogenesis. While progress continues to be made in our understanding of the pathways that control the apical (top) geometry of epithelial cells, we know comparatively little about those that control cell basal (bottom) geometry. To examine this issue, we used the fly ommatidium, which is the basic visual unit of the compound eye. The ommatidium is shaped as a hexagonal prism, and generating this three-dimensional structure requires ommatidial cells to adopt specific apical and basal polygonal geometries. Using this model system, we find that generating cell type-specific basal geometries starts with patterning of the basement membrane, whereby Laminin accumulates at discrete locations across the basal surface of the retina. We show that the Dystroglycan surface receptor promotes this localized Laminin accumulation. Moreover, our results reveal that localized accumulation of Laminin–Dystroglycan induces polarization of Integrin adhesion in ommatidial cells. This underpins cell basal geometry remodeling by anchoring the basal surface of cells to the basement membrane at specific, discrete locations. Altogether, our work shows that patterning of a basement membrane by generating discrete Laminin domains, can direct Integrin adhesion. In the retina, this pathway generates specific basal polygonal geometries to organize a complex multicellular structure in three-dimensions.

## INTRODUCTION

The function of most organs depends upon epithelial tissues. In these tissues, cells coordinate their polarity to generate distinct apical and basal surfaces. This organization underpins the function of epithelial tissues in mediating polarized transport of macromolecules or gas between compartments. At the apical surface, cells adhere to one another through apical-lateral junctions, which seal the epithelial barrier [1]. Basally, cells are attached to the basement membrane, which, amongst other functions, provides biomechanical support to tissues [2]. In addition to this apical-basal organization, epithelia are often folded and can bend to generate tubes and cavities, to generate physiological compartments. However, our understanding of the mechanisms and pathways which control the organization of epithelia during development remains incomplete.

Most studies examining this issue tend to make use of embryonic tissues, which consist of a homogenous cell population. Using these relatively simple tissues has proven extremely valuable to reveal fundamental pathways in epithelial patterning and morphogenesis. For example, the relative movement of cells in the plane of a tissue can promote tissue elongation [3-7]. Moreover, changes in cell apical or basal surface area can contribute to inducing tissue curvature, ranging from fold and tube formation [8-15], to tissue invagination [16-19]. In principle, any polygonal geometry can be generated at the apical surface of epithelial cells, depending on a cell’s number of apical junctions (i.e. number of neighbors), and junction length. However, as tissues develop, cells tend to adopt a hexagonal geometry at their apical surface to minimize surface energy [20]. The situation is likely to be more complex in tissues where cells acquire specific shapes as they undergo differentiation. A good example of this is found in the fly eye. In this sensory epithelium, each of the three epithelial cell types that make up the apical surface (lens), remodel their geometry to adopt stereotyped shapes. The combination of programed cell death, preferential adhesion between different cell types, and actomyosin regulations contributes to determining these different apical geometries [21-30].

Like for the apical surface, cells can adopt polygonal geometries at their basal surface [7, 31, 32]. However, which pathways control these geometries is not clear. Basally, cells are anchored to the basement membrane through Integrins. These surface receptors are heterodimers consisting of an α and β chain, which can bind to extracellular matrix components of the basement membrane. Integrin ligands include Laminins [33, 34], and Collagen IV [35, 36], which are both large trimeric proteins. Integrin binding to these ECM components connects the basement membrane to the F-actin and microtubule cytoskeleton through multiple adaptor proteins such as Talin and ILK [37-39]. In developing epithelia, Integrins have been shown to regulate the basal area of cells. This is the case in the follicular epithelium for example, where they organize the basal F-actin cytoskeleton [40, 41]. In this epithelium, another basal surface receptor, Dystroglycan (DG), also contributes to organizing the basal F-actin cytoskeleton [42, 43]. DG can bind to Laminins [44] and its function in organizing basal F-actin in follicular cells depends upon formation of Laminin fibrils in the underlying basement membrane [42]. Interestingly, work using cultured mouse embryonic stem cells has also revealed a requirement for DG in organizing Laminin-1 within the extracellular matrix, during embryoid body formation [45]. Thus, Laminin organization and DG are intimately linked in epithelial morphogenesis and this relationship appears to be conserved through evolution. Whether and how ECM regulation, DG and Integrins might control the basal geometry of cells during epithelial morphogenesis has not been investigated in any system.

To examine this issue, we made use of the genetically amenable *Drosophila* retina. The retina is made of 750 basic visual units called ommatidia. Each ommatidium is shaped as a hexagonal prism, and within this three-dimensional structure, different cell-types adopt specific basal polygonal geometries [46]. Using this model system, we show that establishing cell basal polygonal geometry involves a switch in Integrin localization, from a basal cluster to polarized localization. We find that this Integrin polarization is directed by LamininA/B1, which accumulate at discrete locations across the ECM of the basement membrane that lines the retinal epithelium. Furthermore, our results indicate that the DG-Dystrophin pathway is required to promote this localized Laminin accumulation. We conclude that during epithelial tissue development, patterning of the basement membrane through DG and Laminins can control the basal polygonal geometry of cells, by directing the location of Integrin adhesion to discrete subcellular domains.

## RESULTS

### Retinal cells remodel their basal geometry to shape the ommatidium as a hexagonal prism

To characterize cell basal geometry remodeling in the retina, we made use of confocal tomography and 3D segmentation. During early ommatidial morphogenesis, cells present poorly defined apical and basal geometries (Figure 1A-B, Movie S1-S2). As ommatidial morphogenesis proceeds, cells progressively remodel their apical and basal geometries to shape the ommatidium as a hexagonal prism (Figure 1C-D, Movie S3-S4). Basally, the secondary pigment cells adopt a stereotyped oblong geometry, and the tertiary pigment cells adopt a triangular geometry [46] (Figure 1E). The bristle cell complex, which consist of four cell types, form a triangular shape, which alternates with the tertiary pigment cells, in between secondary pigment cells (Figure 1C and 1D). At the tissue level, the secondary and tertiary pigment cells which contribute to two and three neighboring ommatidia, respectively, generate a supracellular lattice that connects ommatidia across the basal surface of the epithelium (Figure 1C and Supplementary Figure 1).

**Figure 1:**
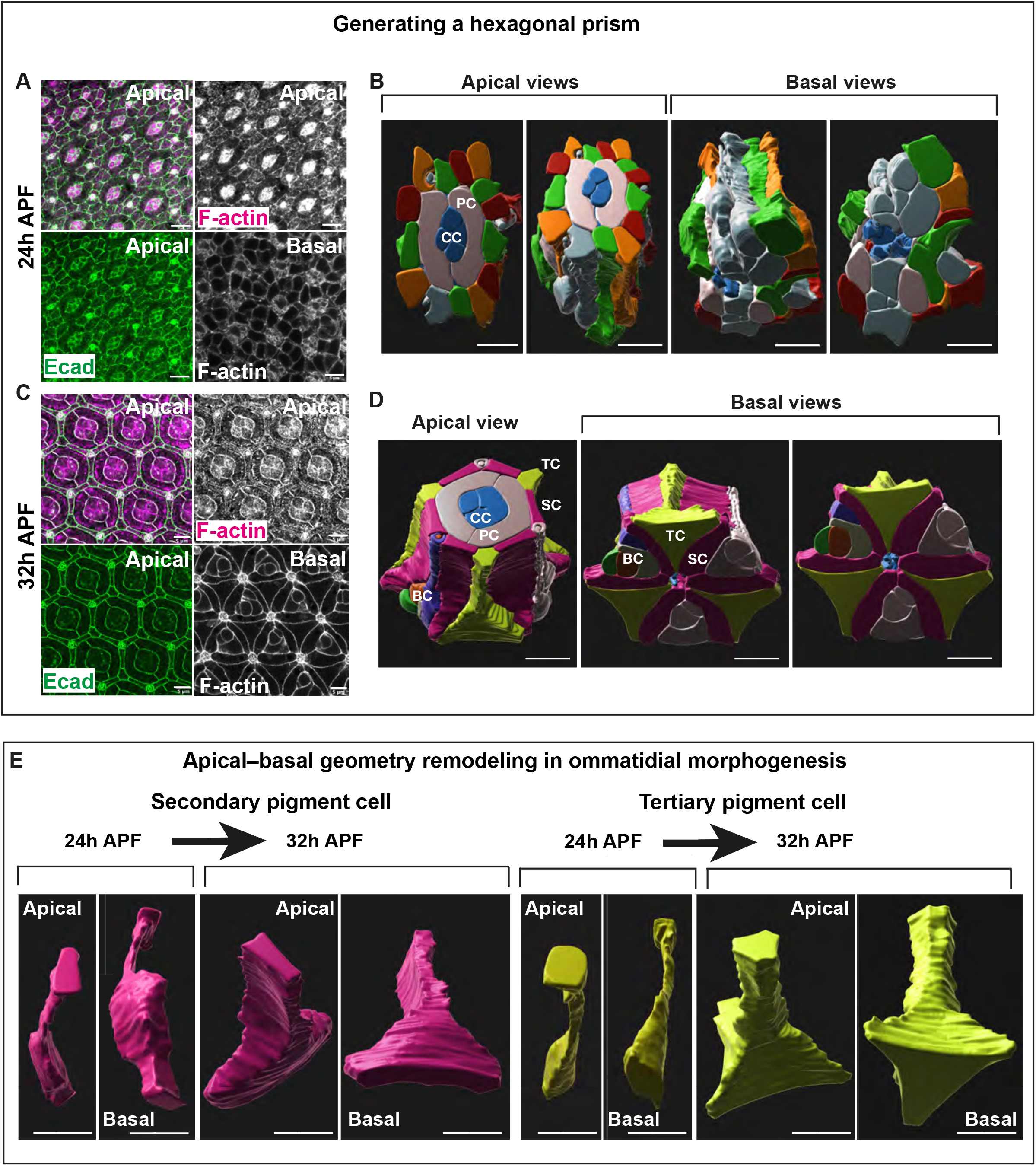
Retinal cells acquire specific basal polygonal geometries in morphogenesis. **(A)** Confocal images of the apical and basal surfaces of an early pupal retina (24h APF). F-actin (magenta) and Ecadherin::GFP labels the apical Adherens Junction (AJ) (green). **(B)** 3D segmentation and rendering of an early ommatidium (24h APF), showing apical and basal side views. The cone cells (CC), at the center of each ommatidium are labelled in blue. Primary pigment cells (PC) are labeled in light grey. The other cells are labelled with random colors to help track them along the apical-basal axis. This shows that at this stage of development, cells do not always share the same neighbors from apical to basal. **(C)** Confocal images of the apical and basal surface of the same patterned pupal retina (32h APF). F-actin (magenta) and Ecadherin::GFP labels the AJ (green). **(D)** 3D segmentation and rendering of a patterned ommatidium (32h APF) showing apical and basal side views. Cone cells (CC, blue), primary pigment cells (PC, grey), secondary pigment cells (SP, magenta), tertiary pigment cells (TC, yellow) and bristle cell complex, which consist of four cells (BC). Scale bars: 5 μm. **(E)** Individual secondary (magenta) and tertiary (yellow) before (24h APF) and after (32h APF) geometry remodeling. Scale bars: 5μm

### Integrins become polarized during cell basal geometry remodeling

To understand how cell basal geometry remodeling is induced, we sought to identify gene requirements in this process. The Integrin *βPS* subunit (Myspheroid, Mys) has been shown to be required to maintain surface integrity late in retinal development, as the tissue surface undergoes basal contraction [46]. Here, we asked whether Integrin adhesion is also required for cell basal geometry remodeling, which precedes contraction. Monitoring the expression of the βPS/Mys subunit revealed that before cell basal geometry remodeling commences, Integrins are mostly concentrated in a cluster at the basal surface of cells (Figure 2A-B). As the interommatidial cells remodel their basal geometry at around 32h APF, βPS/Mys becomes polarized within each ommatidial cell, in the plane of the basal surface of the tissue. This polarization leads to the accumulation of βPS/Mys around the center of the ommatidium (Figure 2C). This generates what has been referred to before as a grommet structure, which anchors the interommatidial cells around the basal feet of the cone cells and the afferent photoreceptor axons [46] (Figure 2C-D). At the more distal region of the grommet, βPS/Mys accumulates in the interommatidial cell feet, surrounding a second population of βPS/Mys that is observed in puncta associated with the basal feet of the cone cells (Figure 2D). More proximally, βPS/Mys accumulation is observed only within the interommatidial cell feet, forming a supracellular ring (i.e. grommet) (Figure2D). Further examination of the basal surface of the cone cells revealed that they present two Integrin αPS subunits, αPS1 (Multiple edematous wing; Mew) and αPS2 (Inflated/If), together with βPS/Mys (Figure 2E and Supplementary Figure 2). In contrast, within the grommet, the interommatidial cells only present αPS1/Mew together with βPS/Mys (Figure 2F and Supplementary Figure 2). Therefore, different retinal cell types present different combinations of α and β Integrin subunits. To confirm that the Integrin staining at the grommet is contributed by the interommatidial cells, we generated mosaic tissue for *talin* (*rhea*) RNAi, an essential component of the Integrin adhesion complex. In secondary and tertiary pigment cells expressing talin RNAi, βPS/Mys localization at the grommet was lost, demonstrating that βPS/Mys accumulation at the grommets results from the polarization of Integrins in the interommatidial cells (Supplementary Figure 3).

**Figure 2:**
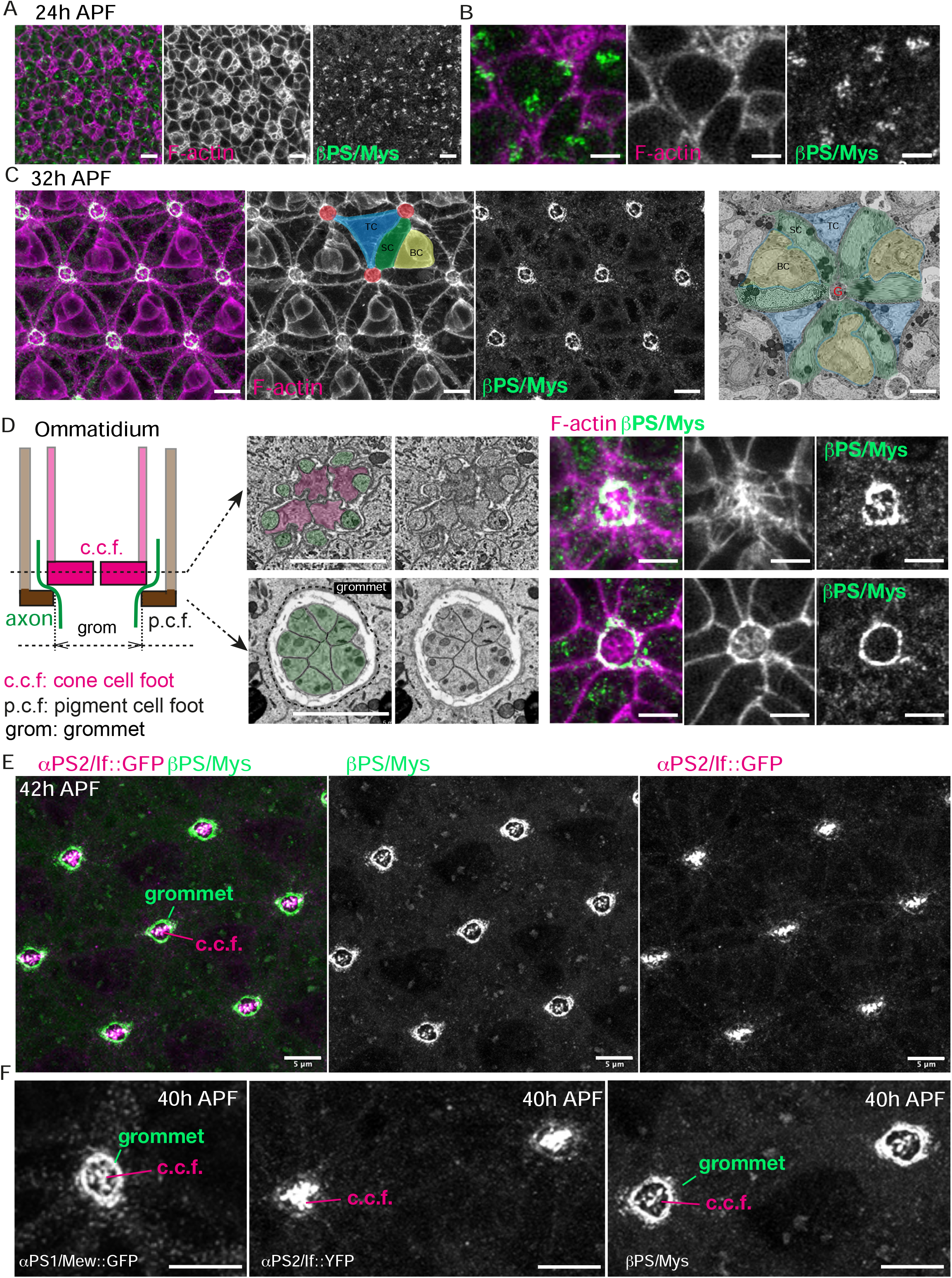
Integrins become polarized as cells remodel their basal geometry. **(A)** Basal surface of a 24h APF retina, before cell basal geometry remodeling has occurred. F-actin (magenta) and Mys/μPS (green). **(B)** Higher magnification of the basal surface of retinal cells prior to geometry remodeling. **(C)** Basal surface of a 32h APF retina, as cells have acquired their basal geometry, F-actin (magenta) and μPS/Mys (green). One secondary (SC, green) and one tertiary (TC, blue) pigment cell is highlighted to show how these cells connect multiple neighboring ommatidia. The center of the ommatidia that they connect are labelled in red. A bristle cell complex (BC) is highlighted in yellow. One electron micrograph of a basal section of the retina is shown, centered on one ommatidium. SC: secondary pigment cells (green), TC: tertiary pigment cells (blue), BC: Bristle cell complex (yellow). Grommet structure (G) around the afferent photoreceptor axons (a). **(D)** Simplified (2D) drawing of a sagittal section of the ommatidium, showing two of the four cone cell feet (c.c.f, magenta) and two of the 6 interommatidial cell feet (I.c.f, brown). The afferent photoreceptors axons (green) run at the periphery of the c.c.f and exit the ommatidium in between the I.c.f. The grommet structure is delineated by the plasma membrane of the pigment cells which surround the c.c.f and axons. The electron micrographs show cross sections cutting through the feet of the cone cells (magenta) and axons (green) (distal section) and through the axons only, below the feet of the cone cells (proximal section). The confocal staining shows optical cross sections at this location, showing that βPS/Mys accumulates at the feet of the cone cells and at the grommet. **(E)** Basal surface of a 42h APF retina αPS2/If::GFP (magenta) and βPS/Mys (green). The αPS2/If::GFP panel shows how this subunit is detected at the cone cell feet, and not at the grommet. **(F)** Projection of confocal section spanning the c.c.f and grommet focal plans. Note how the cone cells express both αPS1/Mew and αPS2/If, while the pigment cells delineating the grommet only express αPS1/Mew. Scale bars **(A**,**C**,**E**,**F):** 5μm and **(B, D):** 2μm

### Integrins are required for cell basal geometry remodeling

Next, to examine the requirement of Integrin adhesion in basal geometry remodeling, we used RNAi. Firstly, we sought to quantify the basal geometry of wildtype interommatidial cells. For this we used automated segmentation with manual correction [47] to measure cell shape parameters at the basal surface of wildtype retina. We focused our analysis on the secondary and tertiary pigment cells, and we used a principal component analysis to describe their basal geometry. This approach allows for a clear distinction between the secondary and tertiary pigment cells (Figure 3A). Decreasing the expression of *talin*, we found that basal surface organization was highly perturbed when compared to wildtype (Figure 3B), and the shape of the secondary and tertiary pigment cells became undistinguishable. To complement these experiments, we expressed a version of Mys/βPS that does not allow for adhesion but that retains signaling capability (Mys^DN^) [48]. Similar to *talin* RNAi, expressing the Mys^DN^ transgene interfered with basal surface organization, as the shape of the secondary and tertiary pigment cells became undistinguishable (Figure 3C). Further, generating *mys*^*1*^ mutant clones led to a cell autonomous disruption of cell basal geometry (Figure 3D). From these experiments, we conclude that Integrin adhesion is required for cell basal geometry remodeling during retinal morphogenesis.

**Figure 3:**
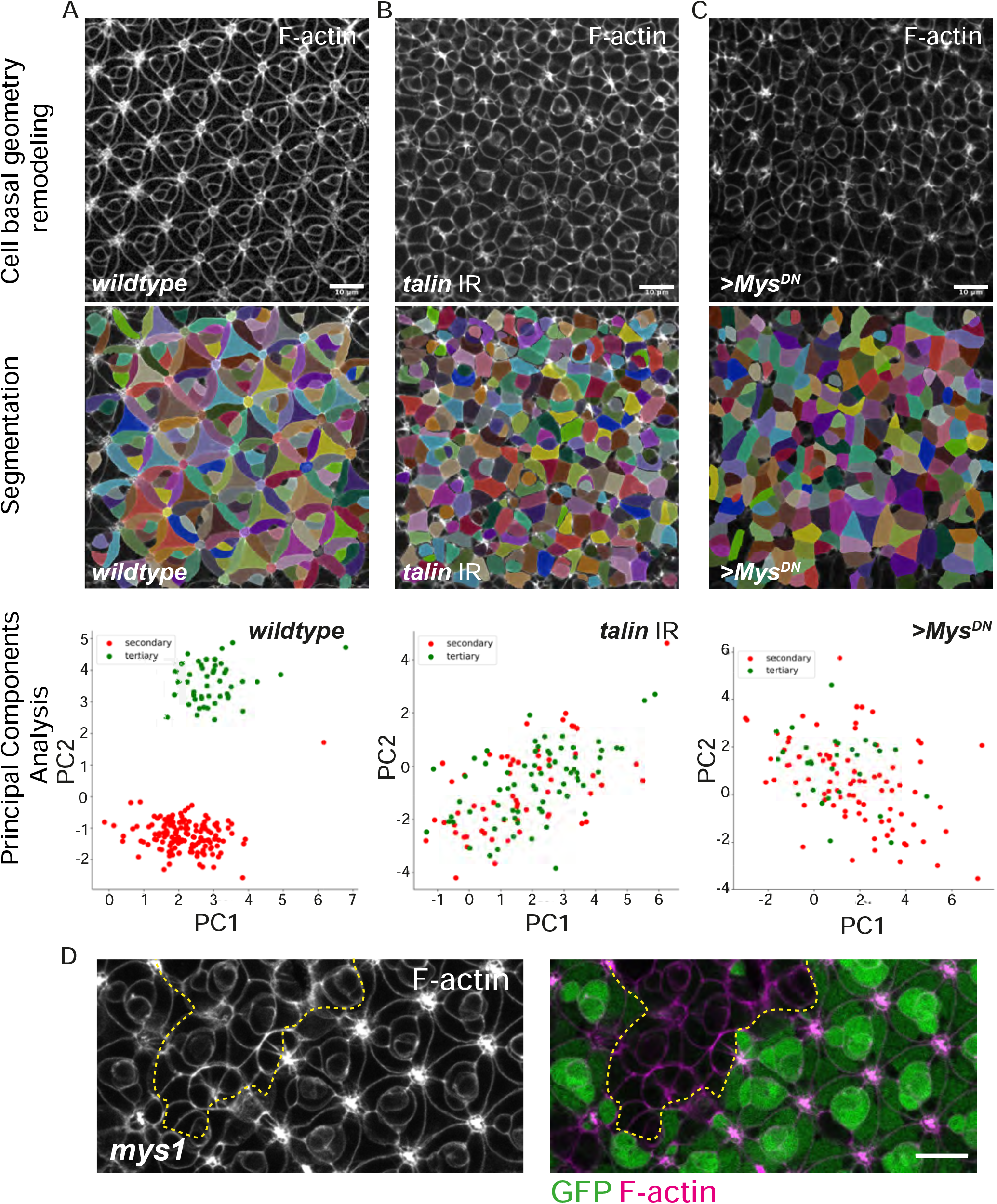
Integrin adhesion is required for cell basal geometry remodeling. **(A)** Confocal section of a wildtype patterned retinal basal surface at 42h APF, segmented using napari and processed through a principal component analysis allowing us to distinguish the basal geometry of the secondary (red) and tertiary (green) pigment cells. **(B)** Confocal section of a *talin* RNAi retinal basal surface at 42h APF, segmented using napari and processed through a principal component analysis allowing us to distinguish the basal geometry of the secondary (red) and tertiary (green) pigment cells. **(C)** Confocal section of a *Mys*^*DN*^ retinal basal surface at 42h APF, segmented using napari and processed through a principal component analysis allowing us to distinguish basal geometry of the secondary (red) and tertiary (green) pigment cells. **(D)** Basal optical section of a *mys*^*1*^ mutant clone (tissue lacking GFP; circled using a yellow dashed line). F-actin (magenta) GFP (green). Scale bars: 10μm.

### Basal geometry remodeling starts with patterning of the ECM through localized LamininA/B1 accumulation

Basement membrane organization influences morphogenesis. This prompted us to examine the relationship between the ECM, Integrin adhesion and cell basal geometry remodeling. To this end, we examined the localization and requirement of the Laminin A and B1 subunits, Perlecan/Trol, Collagen IV/Viking, the glycoprotein Nidogen (Entactin/Ndg), and the secreted glycoprotein protein-acidic-cysteine-rich (Sparc), which are all components of the basement membrane [2]. Using available functional GFP protein traps [49, 50], we found that as early as 20h APF, 12h before the αPS1/Mew – βPS/Mys Integrin receptor is polarized in the interommatidial cells, both the laminin α (LanA) and β (LanB1) chains, accumulate at the center of the ommatidium, in a pattern resembling the grommet structure (Figure 4A and Supplementary Figure 4). At this developmental stage, Collagen IV, Nidogen, and Perlecan are not enriched in the same manner and instead, are distributed across the basal surface of the retina (Figure 4B and Supplementary Figure 5). LamininA/B1 accumulation at the presumptive grommet precedes Integrin accumulation at this location. It suggests that localized Laminin might control Integrin localization in the interommatidial cells.

**Figure 4:**
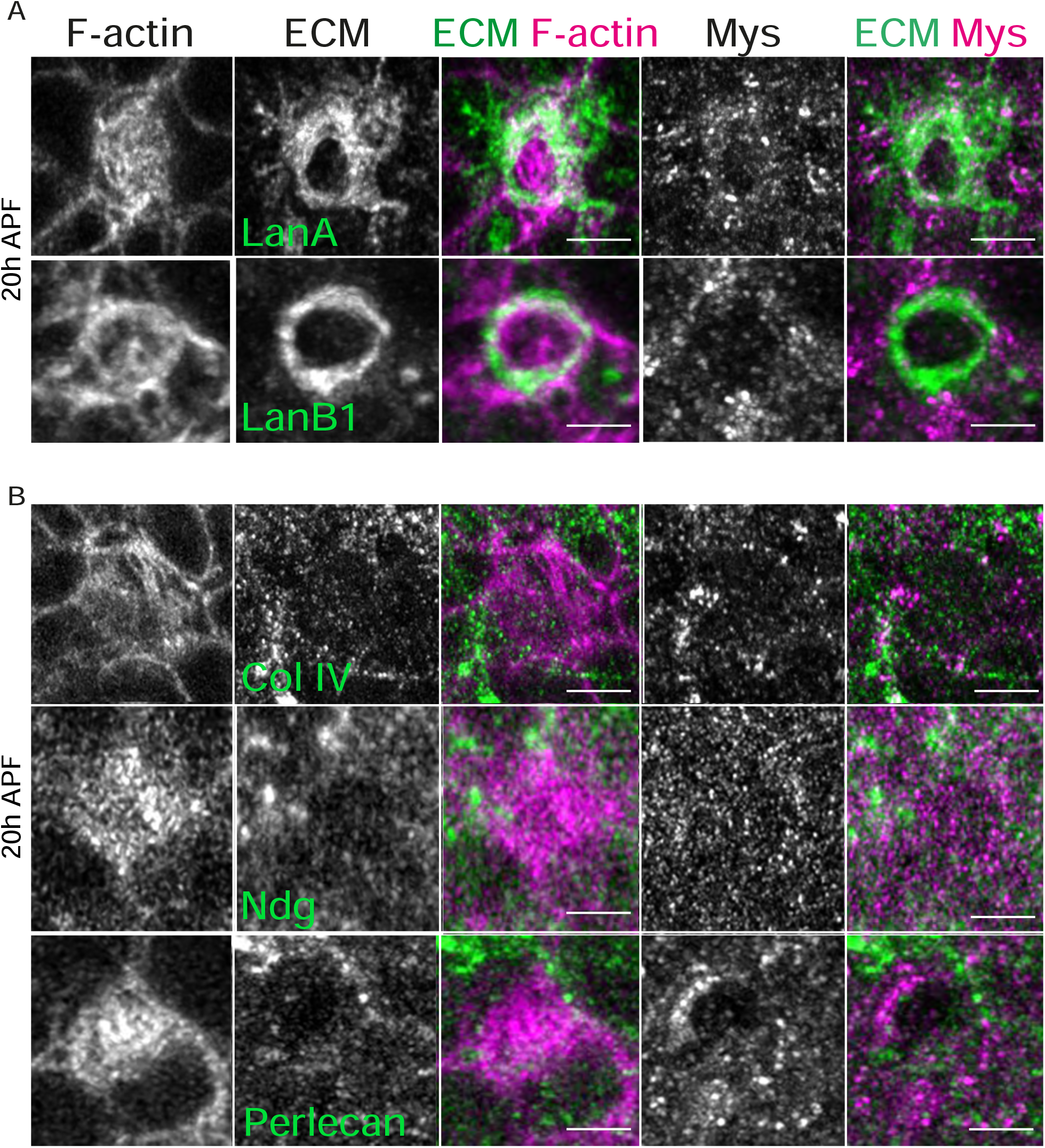
LamininA/B1 accumulate at the grommet before Integrins. **(A-B)** Basal surface of a retina imaged at 20h APF, before the onset of cell basal geometry remodeling. **(A)** The α-laminin subunit LanA and β-laminin subunit LanB1 accumulate at the presumptive grommet. F-actin (magenta), LanA and LanB (green), βPS/Mys (magenta). **(B)** Collagen IV, Ndg and Perlecan show a punctate distribution at the ECM, with no enrichment around the grommet. F-actin (magenta), Col IV, Ndg, Perlecan (green), Mys (magenta). Scale bars: 2μm.

Examining the distribution of the ECM components at later stage of retinal development, when βPS/Mys is concentrated at the grommet, revealed that in addition to the LamininA/B1 subunits, Collagen IV, Nidogen, and Perlecan all become enriched at this location (Figure 5). Interestingly, these components accumulate at different locations. In distal sections, LamininA and B1 localize around the cone cell feet and around the axons of the photoreceptors, and in more proximal sections, at the grommet. In distal sections, Collagen IV and Perlecan are localized around the axons, but not in between the cone cell feet (Figure 5E). In more proximal sections, these two factors are localized at the grommet. Nidogen accumulates in sections below the cone cell feet and proximally, at the grommet. Examining Sparc expression revealed this protein localizes in punctate structures within the photoreceptor soma and axon (Supplementary Figure 6). These specific patterns of expression for LamininA/B1, Collagen IV, Perlecan, Nidogen and Sparc are summarized in (Figure 5E).

**Figure 5:**
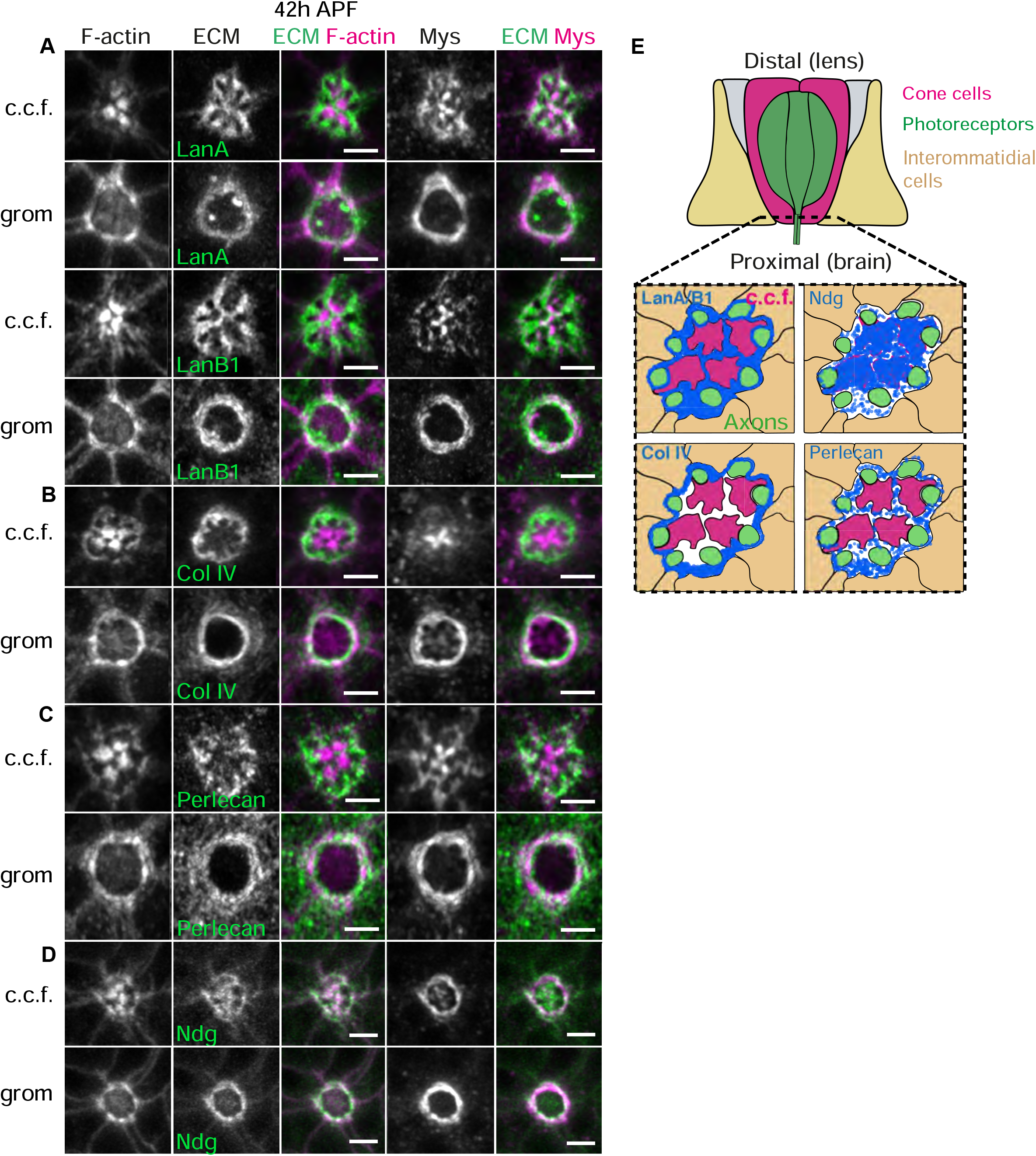
LamininA/B1, Col IV, Perlecan and Ndg distribute differentially at the base of the ommatidium. **(A-D)** Basal sections at the center of one ommatidium, at 42h APF, when cell basal geometry remodeling is completed. **(A)** LanA and LanB1 localization at a distal section, where the feet of the cone cells contact the ECM, and at a proximal section, bellow the feet of the cone cells (c.c.f.), to show the grommet (grom) structure encircling the photoreceptor axons. LanA/B (green) and F-actin (magenta). Distally, LanA/B1 localize around the axons and in between the cone cell feet. Proximally, LanA/B localize at the grommet. **(B)** Collagen IV localization at a distal section, where the feet of the cone cells contact the ECM, and at a proximal section, bellow the feet of the cone cells, to show the grommet. Col IV (green) and F-actin (magenta). Distally Col IV localizes around the axons but is not enriched in between the cone cell feet. Proximally, Col IV localizes like LanA/B1, at the grommet. **(C)** Perlecan localization at a distal section, where the feet of the cone cells contact the ECM, and at a proximal section, bellow the feet of the cone cells, to show the grommet. Perlecan (green) and F-actin (magenta). Distally, Perlecan localizes in punctate structures, around the axons but not in between the cone cell feet. Proximally Perlecan localizes at the grommet. **(D)** Distally, Ndg localizes immediately below the cone cell feet. Proximally, Ndg localizes at the grommet. **(E)** Drawing recapitulating the pattern of expression of LanA/B1, Col IV, Perlecan and Ndg in the distal part of the ommatidium center. Scale bars **(A, C, E)**; 5μm **(B**,**D**,**F)** 2μm

### LamininB2 is required for Integrin polarization during cell basal geometry remodeling

Next, to test the idea that localized Laminin accumulation induces interommatidial cell basal geometry remodeling by recruiting Integrins, we sought to perturb the expression of this ECM component, while monitoring βPS/Mys localization. For this, we used RNAi targeting LanB2, which is common to all possible laminin heterotrimers in *Drosophila*. Consistent with our model that Laminin recruits Integrins at the grommet, we found that LanB2 RNAi leads to defects in βPS/Mys Integrin localization. βPS/Mys fails to accumulate at the grommet, and instead is distributed at the basal plasma membrane into punctate domains (Figure 6A-D). In addition, these perturbation experiments affected cell basal geometry remodeling (Figure 6A, 6C). Consistent with ECM regulation being important for cell basal geometry remodeling, we found that degrading the ECM by expressing Matrix metalloproteases in retinal cells leads to a failure in βPS/Mys localization at the grommet and prevents cell basal geometry remodeling (Figure 6E-F). However, inhibiting the expression of Collagen IV, Ndg, Perlecan and Sparc individually, by expressing RNAi against these genes in all retinal cells, did not lead to defects in βPS/Mys localization. In these genotypes, cells remodeled their basal geometry normally (data not shown).

**Figure 6:**
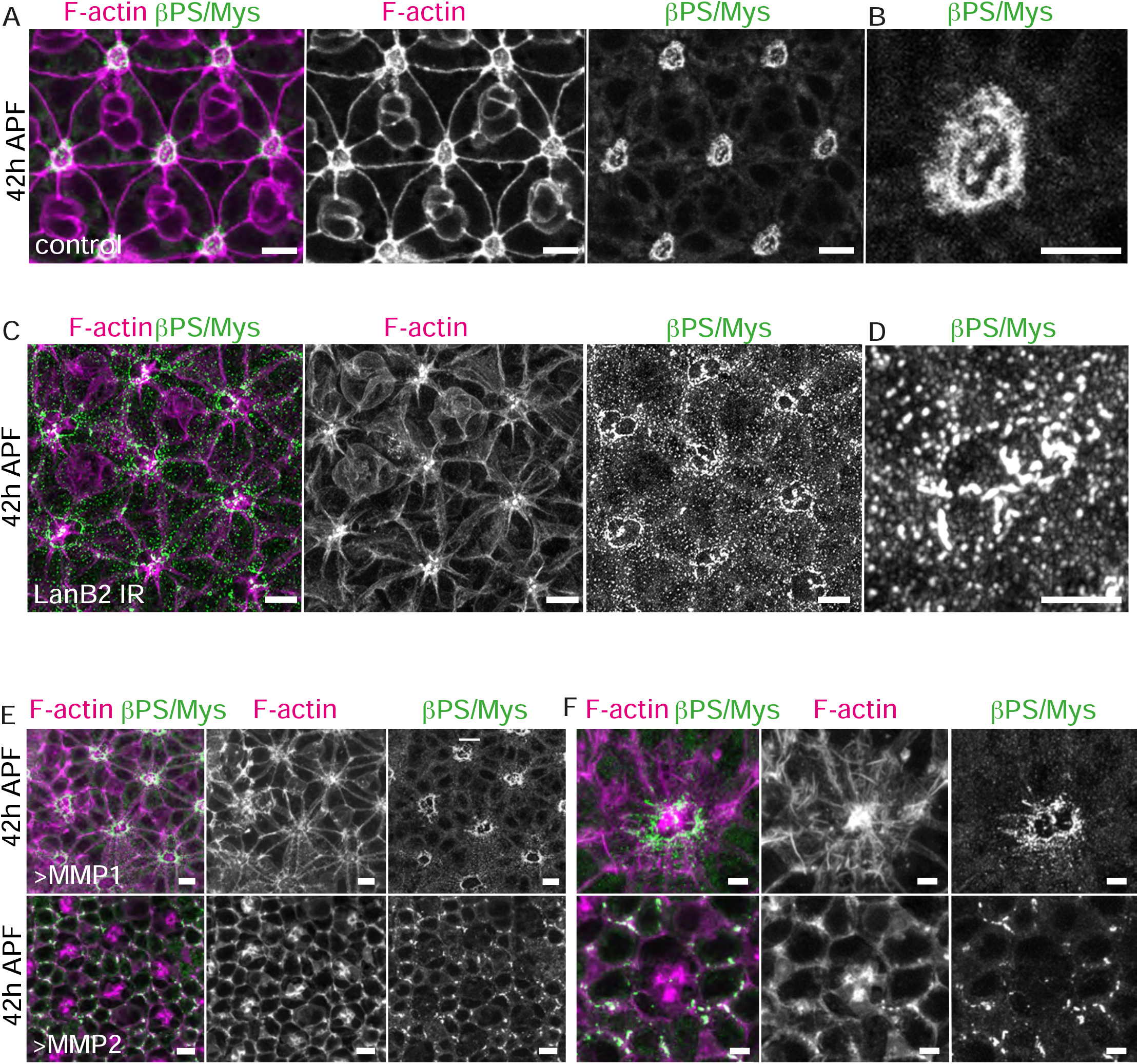
LamininB2 is required for cell basal geometry remodeling. **(A)** Confocal images showing the basal surface of a 42h APF wildtype retina, and **(B)** a close-up onto the center of the ommatidium, where the grommet is located. F-actin (magenta) and βPS/Mys (green). **(C)** Confocal images showing the basal surface of a 42h APF retina expressing LanB2 RNAi, and **(D)** a close-up onto the center of the ommatidium, where the grommet is located. F-actin (magenta) and Mys (green). **(E)** Mmp1 and Mmp2, and **(F)** a close-up onto the center of the ommatidium, where the grommet is located. F-actin (magenta) and βPS/Mys (green).

### DG promotes localized Laminin accumulation

Our experiments suggest an early role for LamininA/B1 in promoting accumulation of the αPS1/Mew – βPS/Mys Integrin receptor at the grommet. DG is required to organize the ECM in several experimental settings [42, 43, 45, 51], which led us to examine its function in localizing Laminin within the retinal ECM. To examine the expression of DG we made use of a Dystrophin::GFP (Dys::GFP) exon trap transgenic strain [52]. Dys is recruited by DG at the plasma membrane, and thus it is suitable for monitoring DG localization. We found that Dys::GFP accumulates at the presumptive grommet structure approximately 12h before Integrins can be detected at this location (Figure 7A-B). In addition, Dys::GFP also localizes at the Integrin clusters at the basal surface of retinal cells. These results suggest that DG-Dys and LamininA/B1 work together during basal geometry remodeling. Examining the basal surface of retina at 32h APF, as cell basal geometry has been remodeled, showed that Dys::GFP localizes at the grommet and also at the feet of the cone cells (Figure 7C-D). At these locations, the Dys::GFP and βPS/Mys signals do not fully overlap, which is consistent with these two surface receptors assembling distinct adhesion sites (Figure 7D-E).

**Figure 7:**
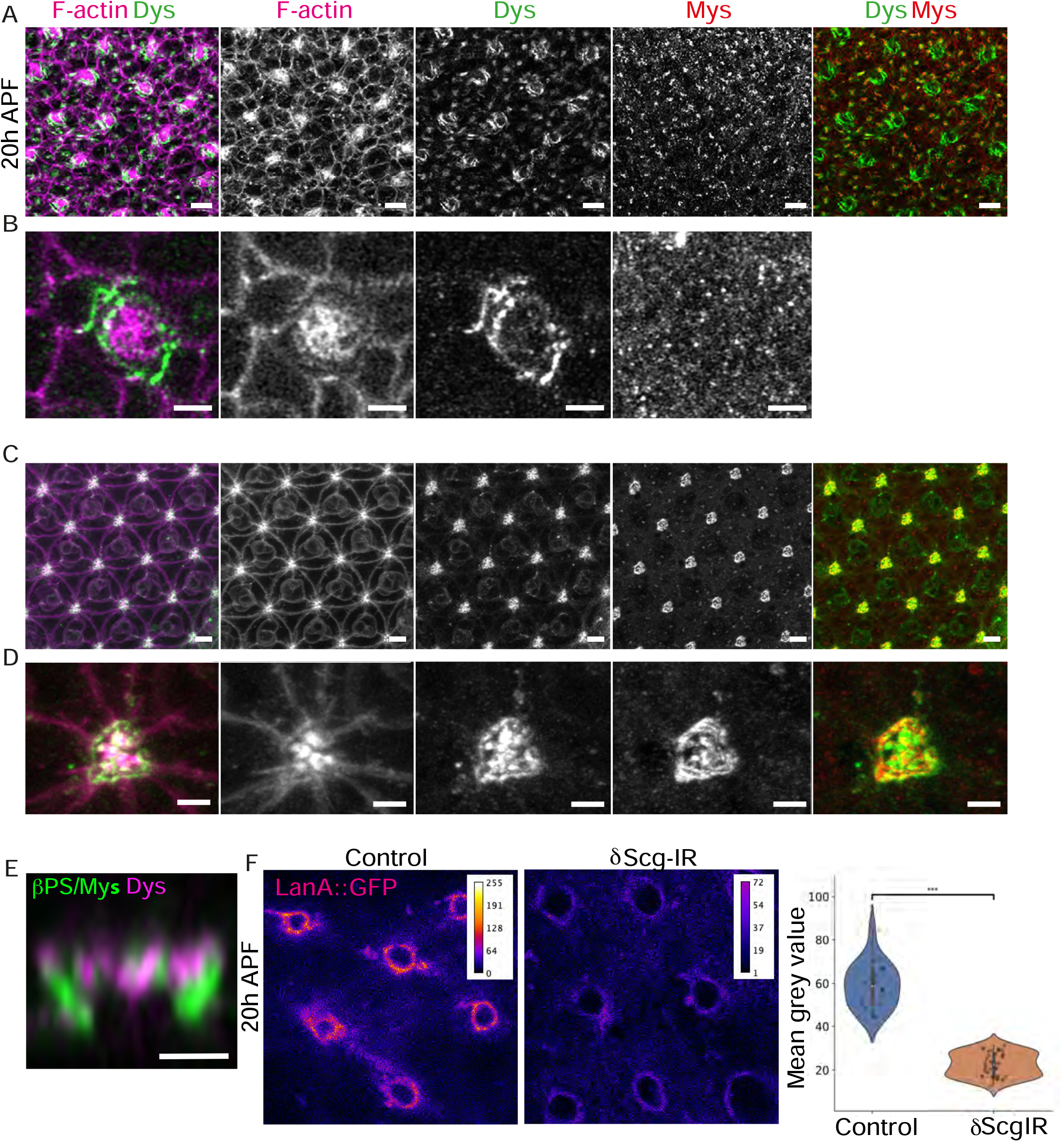
DG is required for localized accumulation of Laminin. **(A)** Confocal images showing a basal retinal surface at 20h APF and a close-up view **(B)** of the presumptive grommet for one ommatidium. F-actin (magenta), Dystrophin::GFP (green), and Mys (red). **(C)** Basal retinal surface at 42h APF and a close-up view **(D)** of a grommet. F-actin (magenta), Dystrophin::GFP (green), Mys (red). **(E)** Sagittal section of a 32h APF grommet showing the relative distribution of βPS/Mys (green) and Dys::GFP (magenta) **(F)** Basal surface of a 20h APF wildtype LanA::GFP, and of a δSarcoglycan RNAi LanA::GFP retina. Quantification of the LanA::GFP signal for these two genotypes. Scale bars: **(A**,**C)** 5μm; **(B**,**D**,**E)** 2μm

The early accumulation of LamininA/B1 (Figure 4) and Dys::GFP (Figure 7A-B) at the presumptive grommet suggest that these two components in cell basal geometry remodeling are linked. To test this idea, we interfered with the DG-Dys pathway using RNAi and asked whether this affected LanA::GFP. We found that inhibiting the expression of the DG cofactor, δSarcoglycan [53] was most effective in inhibiting this pathway in retinal cells. RNAi against δSarcoglycan led to a decrease in LanA::GFP expression at the presumptive grommet at 20h APF (Figure 7F). Conversely, we assessed whether Laminin is required for the early onset of Dys::GFP localization at the presumptive grommet structure. For this, we used RNAi to inhibit the expression of LanB2, the γ-subunit of the Laminin heterotrimer. We found that while we were able to decrease the amount of Dys::GFP at the grommet at around 42h APF, RNAi was not efficient enough to achieve inhibition at the presumptive grommet, at 24h APF (Supplementary Figure 7). This prevented us from conclusively assessing the requirement of Laminin in promoting DG–Dys localization at the presumptive grommet.

### DG is required for Integrin polarization in cell basal geometry remodeling

In our model of cell basal geometry remodeling, DG–Laminin patterns the ECM to direct Integrin adhesion in the plane of the tissue surface. In this model, DG is required for basal cell geometry remodeling. To test this, we used RNAi against DG, Dys and δSarcoglycan. These perturbations led to a failure of βPS/Mys to accumulate at the grommet. Instead, βPS/Mys was localized in punctate structure randomly located at the basal membrane of retinal cells (Figure 8A-B and Supplementary Figure 8). This was accompanied by defects in cell basal geometry remodeling (Figure 8C-E). These results are in good agreement with a model whereby a DG–Laminin– Integrin pathway determines the basal geometry of cells during retinal epithelium morphogenesis.

**Figure 8:**
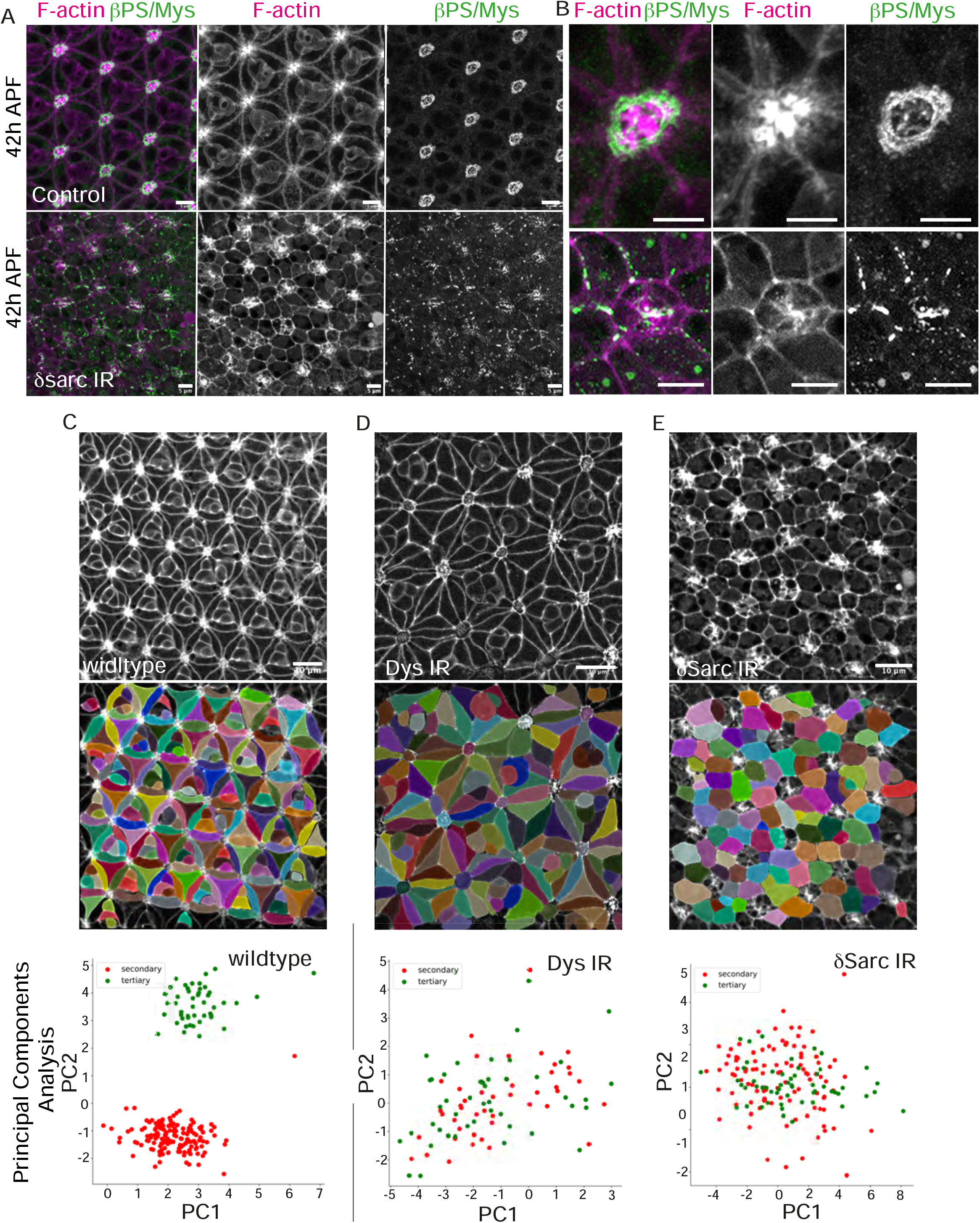
The DG pathway is required for Integrin polarization in cell basal geometry remodeling. **A)** Basal surface of a 42h APF wildtype and δSarcglycan RNAi retina, and **(B)** close-up view of the grommet of these two genotypes. F-actin (magenta) and βPS/Mys::GFP (green). **(C)** Confocal section of a wildtype patterned retinal basal surface at 42h APF, segmented using napari and processed through a principal component analysis allowing us to distinguish basal geometry of the secondary (red) and tertiary (green) pigment cells. **(D)** Confocal section of a *Dys* RNAi retinal basal surface at 42h APF, segmented using napari and processed through a principal component analysis allowing us to distinguish basal geometry of the secondary (red) and tertiary (green) pigment cells. **(E)** Confocal section of a *δsarc* RNAi retinal basal surface at 42h APF, segmented using napari and processed through a principal component analysis allowing to distinguish basal geometry of the secondary (red) and tertiary (green) pigment cells. Scale bars: **(A**,**C)** 5μm; **(B**,**D**,**E)** 2μm

## DISCUSSION

We have used the fly retina as a model system to reveal candidate pathways in controlling the geometry of the basal surface of epithelial cells during morphogenesis. The basal surface of the retina becomes highly organized during development as the interommatidial cells adopt specific apical and basal polygonal geometries. Our results indicate that generating specific polygonal geometries at the basal surface of cells starts with organizing the ECM to establish a pattern of Laminin-rich domains, distributed across the tissue basal surface. We find that establishing this Laminin pattern depends upon the DG pathway. In turn, this DG–Laminin pattern directs recruitment of Integrins, to induce basal geometry remodeling. Thus, we propose that patterning of the ECM, through local Laminin accumulation can determine the basal polygonal geometry of epithelial cells, by controlling the localization of Integrin adhesion. In the retina, failure to establish specific polygonal geometries at the basal pole of cells will compromise eye function, as precise geometry control is key for the function of this organ in detecting object motion.

### DG promotes patterning of the ECM through localized Laminin accumulation

DG is essential for basement membrane formation across model systems. DG knockout mice fail to develop beyond 5.5 embryonic days due to a failure in assembling the extra-embryonic basement membrane, called the Reichert’s membrane [51]. Furthermore, examining DG function during the formation of mouse embryoid bodies shows that this cell surface receptor is required for recruiting and organizing Laminin-1 at the basement membrane [45]. In this system, the authors proposed that DG promotes Laminin-1 recruitment at the plasma membrane, which in turn is thought to promote Integrin binding. In this model system, Integrins re-arrange Laminin-1 into fibrils and plaques [54]. Through this mechanism, DG and Integrins synergize in organizing the ECM. The idea that DG recruits Laminin to enable Integrin adhesion is well supported by work using Caco-2 intestinal epithelial cells [55]. Moreover, this type of relationship between DG, Laminin and Integrins in basement membrane formation, has also been reported in the *Drosophila* oocyte [42, 43, 56]. In this tissue, DG is required to generate Laminin fibrils that line the basal surface of the follicular epithelium, surrounding the oocyte. In addition, DG promotes formation of basal F-actin fibres in the follicular cells, which align with the Laminin fibrils. Integrin adhesion is also required to organize this basal F-actin cytoskeleton, and cells deficient for Integrins present a reduced area when compared to their wildtype counterpart [41, 57]. In this tissue, alignment of the Laminin fibrils and basal F-actin is thought to generate a corset, which mechanically constrains the oocyte to control its axis of elongation. Thus, a combination of DG, Laminin and Integrin adhesion regulates the area of the basal surface of the follicular epithelial cells, and the overall shape of the egg.

In the developing retina, our results show that DG is required in the interommatidial cells to promote Laminin accumulation around the center of the ommatidium, where the feet of the cone cells are attached to the ECM, and where the photoreceptor axons exit the ommatidium. We find that both Laminin and Dystrophin accumulate specifically at this location. Because inhibiting δsarcoglycan leads to a decrease in LanA at the grommet, we propose that DG is involved in recruiting Laminin at this location. Exactly what induces the DG–Laminin interaction at this location is not clear. A simple model could be that the cone cell feet and/or the photoreceptor axons, secrete Laminin. Compatible with this idea, we detect LanA::GFP in punctate structures in the photoreceptor soma and axons. However, similar punctate structures are also present in the interommatidial cells, suggesting that multiple cell types probably produce and deposit this ECM component at the grommet. During retinal tissue morphogenesis, we expect that Laminin promotes DG accumulation at the grommet. However, testing this using RNAi failed to reveal such a requirement. In these experiments, we are expressing RNAi in all retinal cells for a relatively short period of time, and it is likely this technical limitation prevents efficient knock down. Alternatively, the source of early Laminin deposition at the presumptive grommet could be extra-retinal. For example, the glial cells, which enwrap the afferent photoreceptor axons, or circulating hemocytes could be the source of Laminin.

### Controlling Integrin adhesion contributes to determining the basal geometry of cells

By examining the pattern of Integrin expression during retinal tissue development, we found that prior to establishing cell basal polygonal geometries, the βPS/Mys Integrin subunit is localized in clusters at the basal surface of retinal cells. These adhesion sites in retinal cells were previously missed [46]. As cells remodel their basal geometry, the basal Integrin clusters can no longer be detected. Integrin localization becomes polarized in the interommatidial cells as it concentrates at the grommet. Therefore, cell basal geometry remodeling correlates with a switch in Integrin localization. We also note that while the interommatidial cells express αPS1/Mew–βPS/Mys, the cone cells express both αPS1/Mew–βPS/Mys and αPS2/if–βPS/Mys. Thus, different cell types express different αPS subunits. It is not clear why the cone cells express two α-subunits, and it will be interesting to examine the roles that these subunits play in cone cell adhesion to the ECM and what the implications might be when considering Integrin signaling during cone cell morphogenesis.

βPS Integrin and Laminins (LamC1) have previously been implicated in optic cup morphogenesis [11, 14, 58]. In this tissue, βPS Integrin and LamC1 are required for cell basal contraction, which promotes tissue curvature, essential for generating a cup-like structure. Similarly, shaping the developing wing epithelium of *Drosophila* as a dome-like structure also requires Integrin, ECM regulation and basal contractility [59]. As mentioned earlier, Integrins are required to organize F-actin at the basal pole of the follicular epithelial cells, and the loss of this adhesion receptor leads to a reduction in basal surface area [41]. In all these model epithelia, Integrin adhesion organizes the basal actomyosin cytoskeleton and provides anchoring points against which this contractile machinery can pull to generate traction. Integrins have also been implicated in cell basal contraction in the retina, however, earlier work using a thermosensitive loss of function allele of βPS/Mys had ruled out a role this adhesion receptor in cell basal geometry remodeling [46]. Our work using both RNAi and the strong *mys*^*1*^ loss-of-function allele [60], shows a clear requirement for Integrin adhesion in cell basal geometry remodeling. It also supports a model whereby controlling where Integrin adhesion is localized at the basal surface of cells is a mechanism of cell basal geometry control.

Most tissues are lined by a basement membrane containing Laminins, and DG and Integrins mediate basal adhesion in epithelia across species. We therefore envisage that during morphogenesis, the DG–Laminin–Integrin pathway can potentially control the basal polygonal geometry of epithelial cells across tissues and species.

## MATERIAL & METHODS

### Fly strains & genetics

Flies were raised on standard food. The following fly strains were used:

*Ubi-Ecadherin::GFP [61]*

*vkg::GFP and Trol::GFP [62]*

*LanA::GFP, LanB1::GFP*,

*Ndg::GFP and SPARC::GFP [49]*

*Ubi-Myspheroid::GFP [63]*

*UAS-Torso::Mys*^*DN*^ [48]

*Dystrophin::GFP [52]*

*Mew-YFP and If-YFP [64] GMR-Gal4* [65].

*UAS-talin RNAi VDRC 40339, UAS-LanB1 RNAi VDRC 23119, UAS-Vkg RNAi VDRC 106812, UAS-dys RNAi BL55641 and VDRC 106578, UAS-ndg RNA VDRC109625 [66]*

*mys*^*1*^ *FRT19A/Fm7C (BL23862), UAS-talin RNAi BL 33913, UAS-dg RNAi BL34895, UAS-δsarcoglycan RNAi BL55325, UAS-trol RNAi BL38298 and BL29440, UAS-ndg RNA BL62902, UAS-LanB2 RNAi BL62002 [67, 68]*

*UAS-Mmp1 BL58700 and UAS-Mmp2 BL58704 [69]*

RNAi and Mmp1/2 experiments were performed at a range of temperatures, from 25ºC to 29ºC, to modulate the strength of expression through the Gal4/UAS system [70].

### Antibody staining

Retinas of appropriately staged animals were dissected in PBS on ice and fixed in 4% paraformaldehyde for 20mins at room temperature (RT) [71]. Retinas were washed in PBS-Triton 0.3% (PBS-T) then stained with primary antibody in PBS-T for 2hrs at RT or overnight at 4°C. Retinas were washed in PBS-T and then stained with secondary antibodies for 2h at RT or overnight at 4°C. Retinas were mounted in Vectashield (Vectorlabs). The following antibodies were used: rat DCAD2 anti-ECadherin (1:50), deposited to the DSHB by Uemura, T. (DSHB Hybridoma Product DCAD2) [72], mouse CF.6G11 anti-Myspheroid (1:50), deposited to the DSHB by D. Brower, [73], combined with mouse or rat secondary antibodies conjugated to Dy405, Alexa488, Cy3 or Alexa647 (Jackson ImmunoResearch) as appropriate, used at 1:200, and phalloidin-TRITC (Sigma), to visualize F-actin.

### Confocal imaging and image processing

Images of fixed retinas were acquired on a Zeiss 880 Mulitphoton Airyscan confocal microscope, and a Zeiss 900 confocal microscope with Airyscan2. Airyscan images were processed using the default settings offered by Zeiss. All images presented were processed using FIJI [74] and AdobePhotoshop CS4 (Adobe). Graphs were produced in GraphPadPrism 7 (GraphPad) or Python (seaborn and matplotlib). Figures were mounted in Adobe Illustrator CS4 (Adobe). Fiji was used to improve the cell membrane fluorescence signal with background subtraction (Rolling ball radius 50 pixels) and by applying a 3D gaussian blur filter (Sigma 1.5). 3D segmentation was performed on samples stained with Ecadherin and phalloidin to segment cell membranes. Using Imaris 9.1.2, cells were manually segmented. 2D contours were drawn around each cell every two Z slices and surfaces were created to render individual cells in 3D.

### 2D Segmentation

Segmentation was performed on samples stained with phalloidin and Ecadherin to segment cell membranes and to assign cell types respectively. Airyscan confocal images were processed with default Zeiss Airyscan processing parameters. Images were then processed with Napari [47] using the background subtraction filter (Rolling ball radius 50 pixels) and a gaussian blur filter (Sigma/Radius 1.5) to enhance cell membrane signal for 2D segmentation. Segmentation, manual correction, and quantification were performed using Napari. Cells were automatically segmented in 2D using the Cellpose plugin with the cyto1 model. The masks generated from the automatic segmentation were then manually corrected. Cell parameters were calculated from the segmented masks using regionprops from scikit-image.

### Principle component analysis

Principal competent analysis (PCA) was carried out using the Scikit-learn library in Python. The Standard scaler package was used to standardise the data across all metrics before calculating the principal components. The PCA package was then used to perform the PCA. Metrics included in the PCA were as follows: extent, max ferret diameter, major axis length, minor axis length, eccentricity, aspect ratio, roundness, circularity, area, cell shape index, perimeter.

### Electron microscopy

Retinas were prepared for electron microscopy as in [75] but embedded in Epon 812 resin. Serial ultrathin sections were collected on ITO coated coverslips and imaged using a Sense backscattered electron detector in a Gemini 300 SEM with Atlas 5 for array tomography acquisition (Zeiss) – operating at 4.5kV accelerating voltage, with 3kV stage bias.

## FIGURE LEGENDS

**Supplementary Movie 1 and 2:** 3D rendering of a 24h APF ommatidium obtain after airyscan confocal imaging.

**Supplementary Movie 3 and 4:** 3D rendering of a 32h APF ommatidium obtain after airyscan confocal imaging.

**Supplementary Figure 1:**
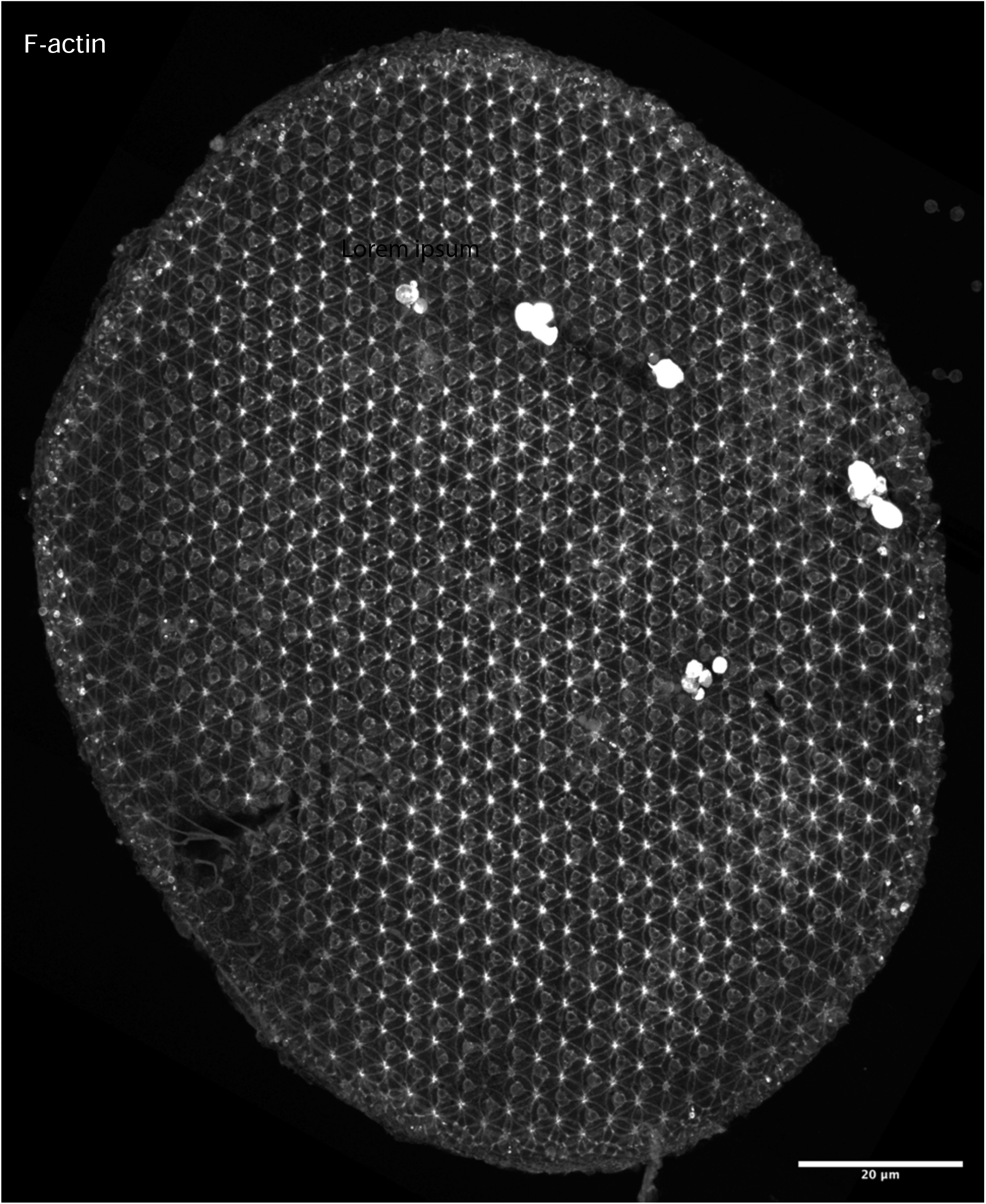
Retinal basal surface organization. F-actin staining revealing cell basal outlines in a 42h APF retina. Scale bar: 20μm.

**Supplementary Figure 2:**
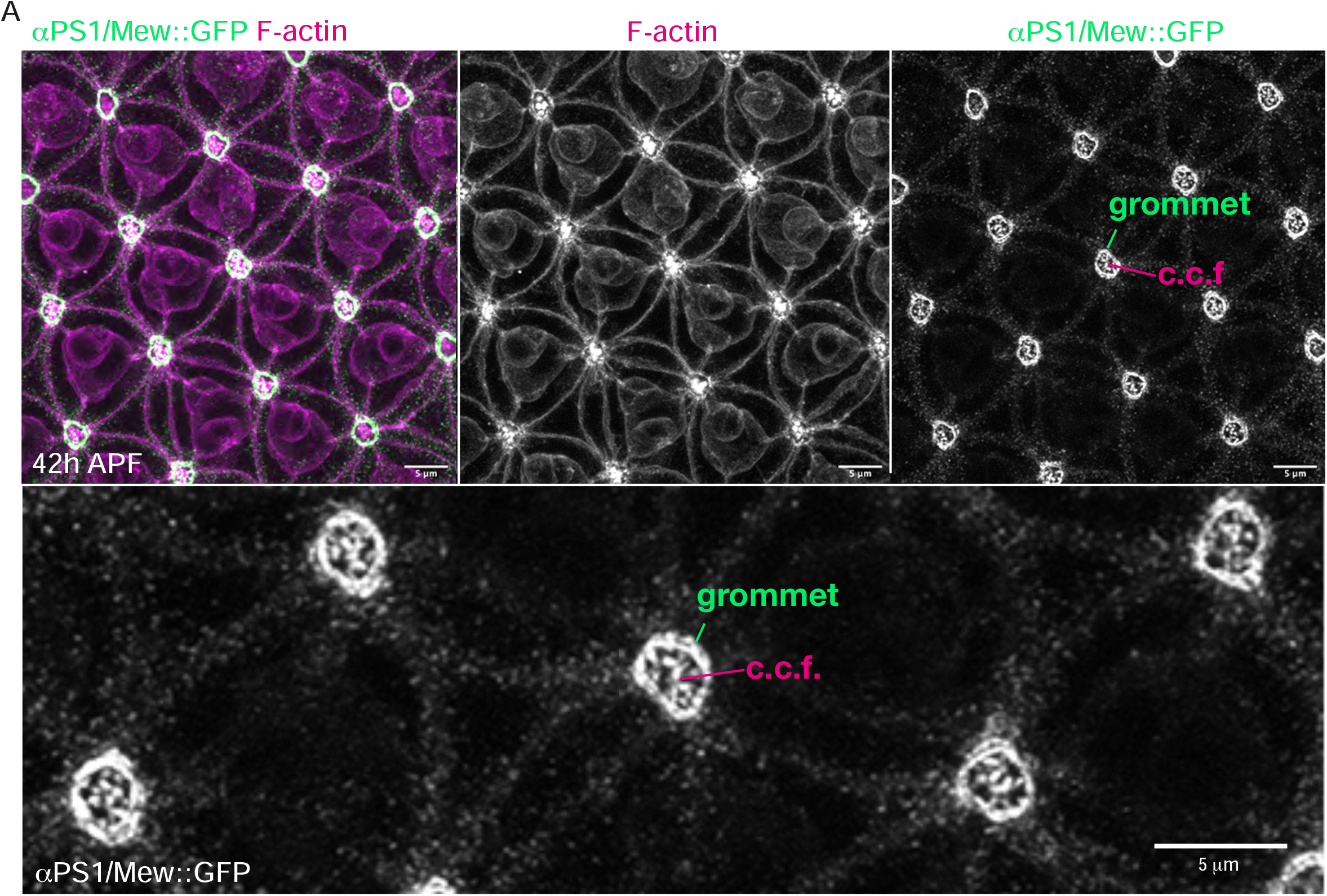
The interommatidial and cone cells express αPS1/Mew. **(A)** Basal surface of a 42h APF retina stained for αPS1/Mew::GFP (green) and F-actin (magenta). Cone cell feet (c.c.f.). The higher magnification shows the αPS1/Mew::GFP localization pattern at the grommet

**Supplementary Figure 3:**
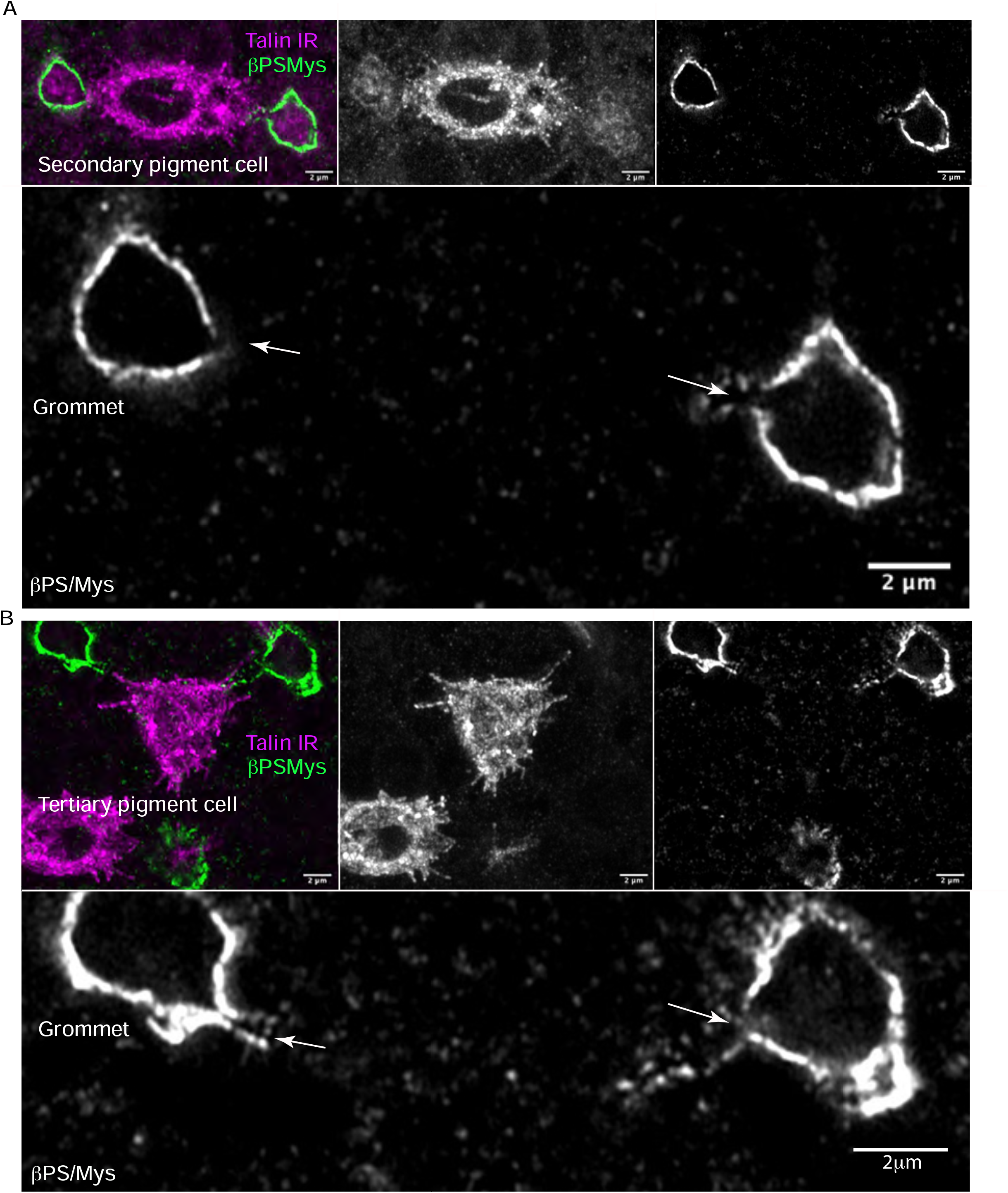
Integrin attachment is polarized in the interommatidial cells. **(A)** 42h APF Secondary pigment cell expressing *talin* RNAi (magenta), surrounded by wildtype cells. βPS/Mys (green). In the close-up view, an arrow points to the lack of βPS/Mys corresponding to where the *talin* deficient cell should contribute Integrins to the grommet **(B)** 42h APF Tertiary pigment cell expressing *talin RNAi* (magenta), surrounded by wildtype cells. GFP (magenta) marks the cell expressing the RNAi. In the close-up view an arrow, points to the lack of βPS/Mys at the location where the cell joins the grommet.

**Supplementary Figure 4:**
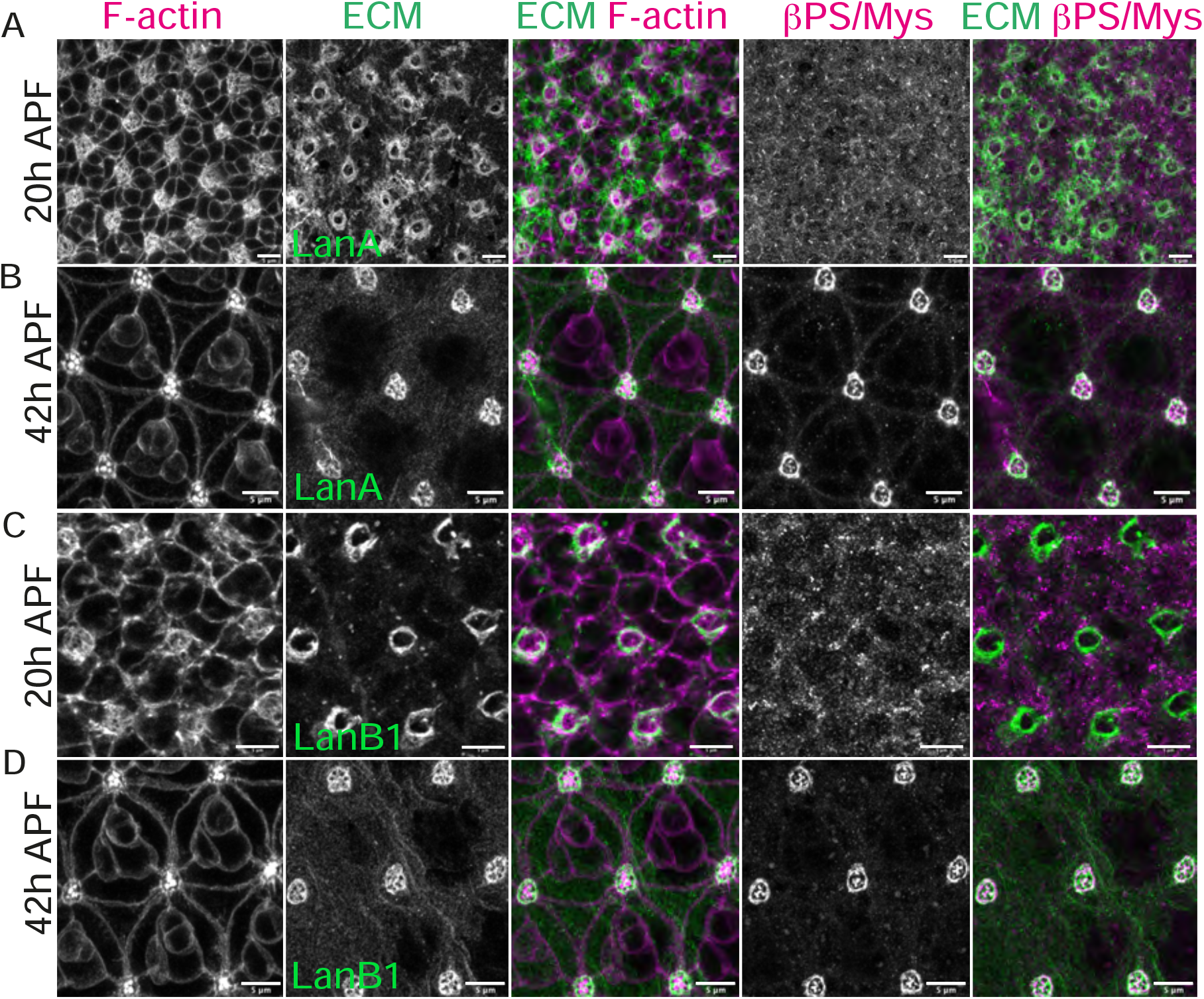
LamininA/B1 localizes at the presumptive grommet prior to Integrins. **(A-B)** Basal surface of a 20h APF and 42h APF retina showing LanA::GFP accumulation at the grommet. **(C-D)** Basal surface of a 20h APF and 42h APF retina showing LanB::GFP accumulation at the grommet. F-actin (magenta), LanA/B (green), Mys (magenta). Scale bars: 5μm

**Supplementary Figure 5:**
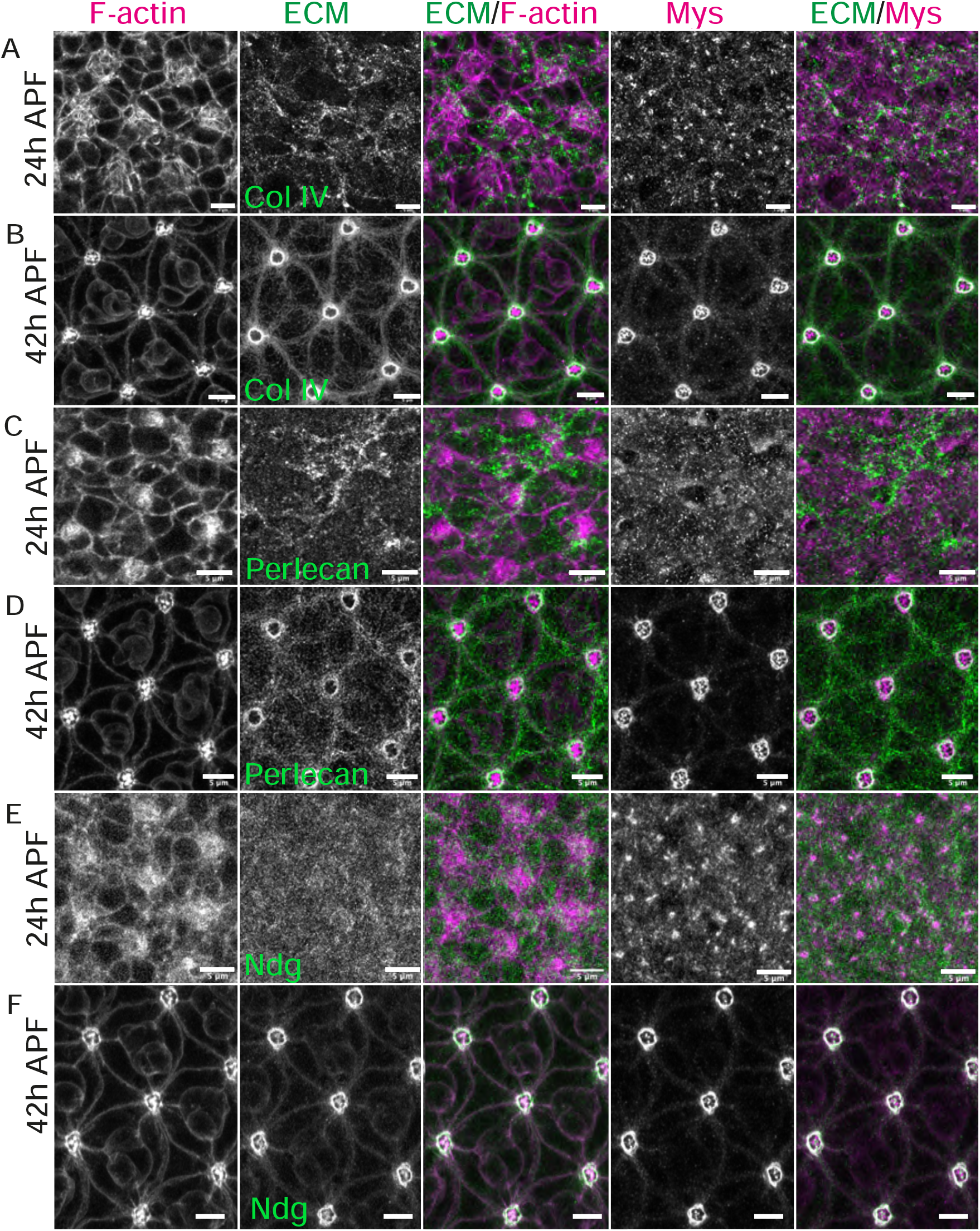
Differential accumulation of ECM factors in basal surface morphogenesis. **(A-F)** Projection of a few contiguous basal sections of retinas at 24h and 42h APF **(A-B)** Col-IV::GFP (green), F-actin (magenta), Mys (magenta). **(C-D)** Perlecan::GFP (green), F-actin (magenta), βPS/Mys (magenta). **(E-F)** Ndg::GFP (green), F-actin (magenta), βPS/Mys (magenta). Scale bars: 5μm

**Supplementary Figure 6:**
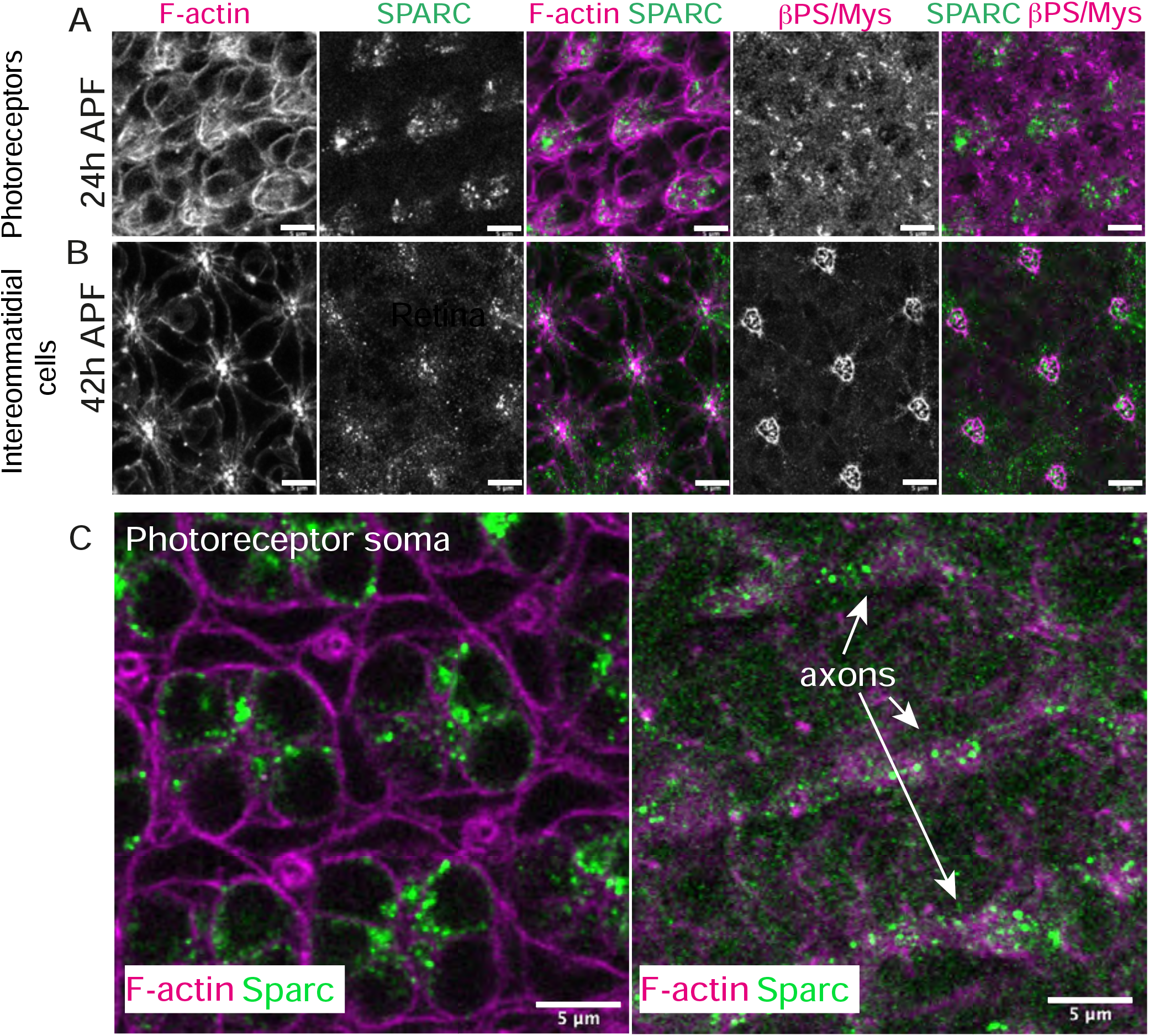
SPARC localizes in the photoreceptor cytosol and axon. **(A)** Projection of contiguous confocal sections taken in the plane of the photoreceptors’ cell body in a 24h APF retina. **(B)** Projection of contiguous confocal sections at the basal surface of a 42h APF retina. **(C)** Close up view on photoreceptors soma in a 42h APF retina. **(D)** Close up view of the basal surface of a 42h APF retina showing thee photoreceptor’s axon bundles running across the tissue surface. F-actin (magenta), SPARC (green) and βPS/Mys (magenta). Scale bars: 5μm

**Supplementary Figure 7:**
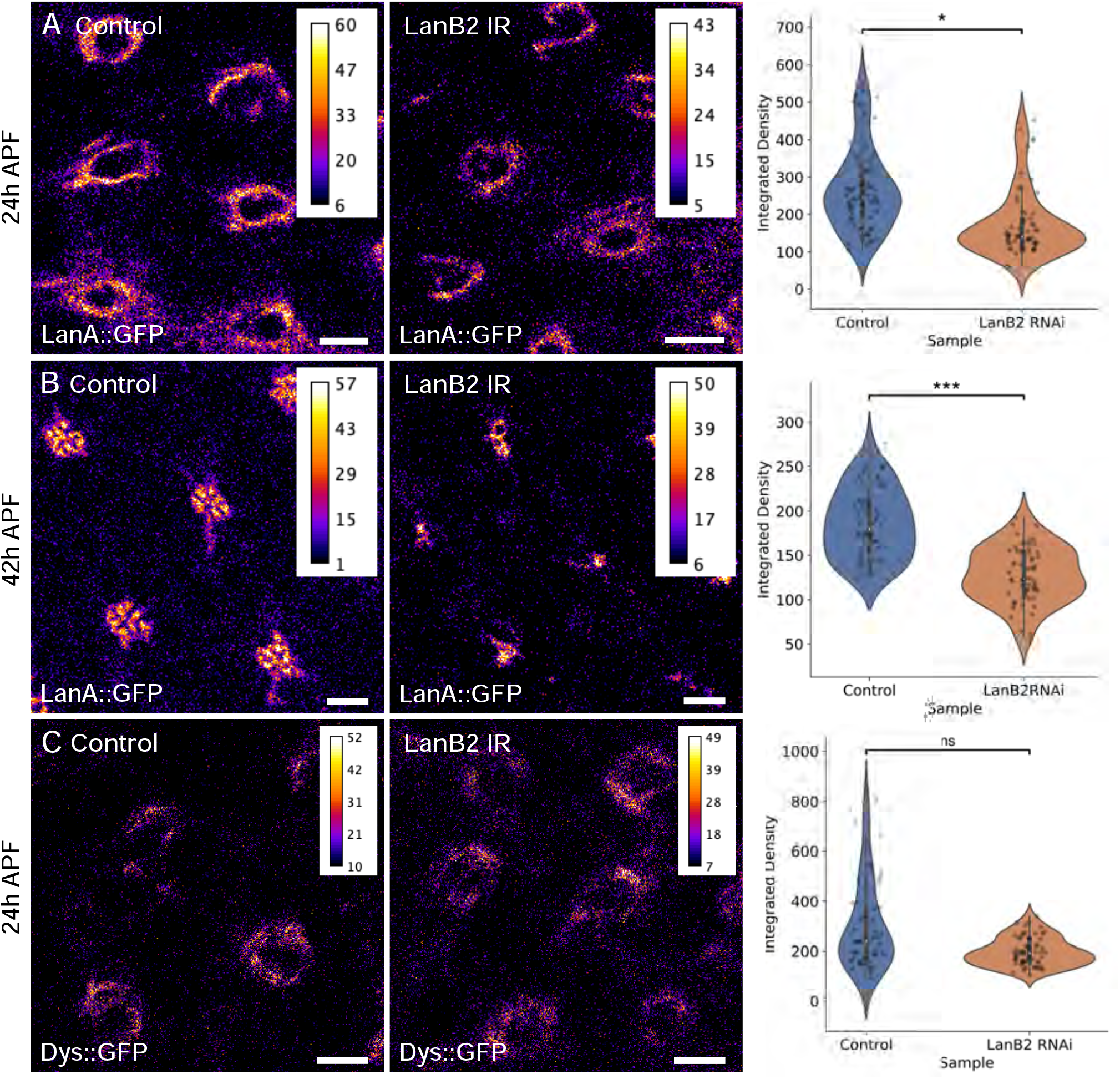
Targeting LanB2 using RNAi does not affect DG localization at the presumptive grommet. **(A)** Basal surface of a wildtype and LanB2 RNAi 24h APF retina. LanA::GFP (fire), and Intensity quantification. **(B)** Basal surface of a wildtype and LanB2 RNAi 42h APF retina. LanA::GFP (fire), and Intensity quantification. **(C)** Basal surface of a wildtype and LanB2 RNAi 24h APF retina. Dys::GFP (fire), and Intensity quantification. Scale bars **(A**,**B**,**C)**: 2μm.

**Supplementary Figure 8:**
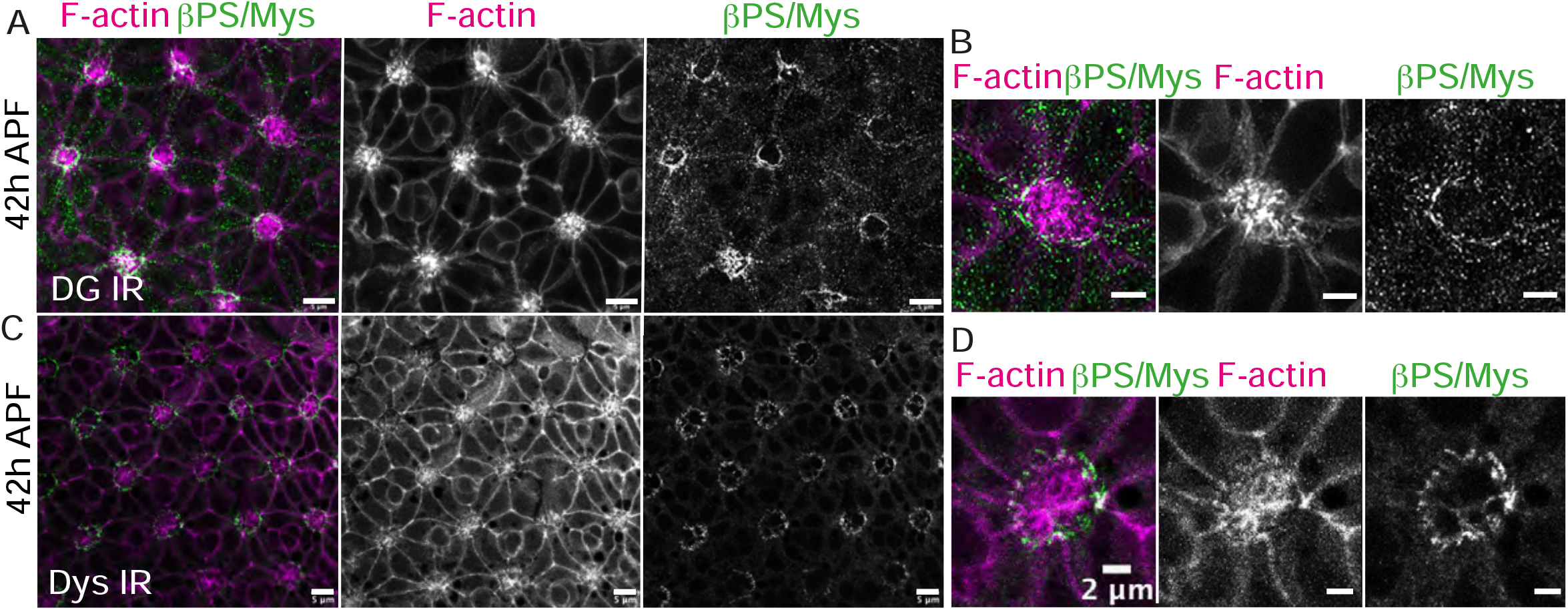
The DG-Dystrophin pathway is required for Integrin polarization. **(A)** Basal surface of a 42h APF retina expressing DG RNAi. F-actin (magenta), βPS/Mys (green). **(B)** Close-up view of a DG RNAi. **(C)** Dys RNAi F-actin (magenta), βPS/Mys (green). **(C)** Basal surface of a 42h APF retina expressing Dys RNAi. F-actin (magenta), βPS/Mys (green). **(D)** Close-up view of a Dys RNAi. Scale bars **(A**,**C)**: 5μm; **(B**,**D)**: 2μm.

## ACKNOWLEDGEMENTS

The authors would like to thank Andrew Vaughan and Ki Hng, who run the LMCB imaging platform and provided training and assistance to RFW and CL for Airyscan confocal imaging. Work in the Pichaud lab is funded by grants from the MRC (MC_UU_12018/3), the BBSRC (BB/R000697), Wellcome (218278/Z/19/Z) and the Royal Society (Award #181274), and benefited from core funding to the LMCB, covering access to microscopy and including support for JJB (MC_12266B). RWF is funded by the MRC (MC_UU_12018/3), and CL is funded through the UCL–Wellcome ‘Optical Biology’ PhD program (218530/Z/19/Z).

## Author contributions

FP and RFW conceived the project. FP wrote the manuscript and prepared the figures, together with CL and RFW. FP and RFW supervised CL’s work. FP, RFW and CL designed the experiments, genetic crosses, and immunostaining/imaging. RFW and CL performed the experiments and analyzed the corresponding data with assistance from FP. JB cut the retinas, prepared by RFW, and imaged the sections using electron microscopy.

